# Cell culture model system utilizing engineered A549 cells to express high levels of ACE2 and TMPRSS2 for investigating SARS-CoV-2 infection and antivirals

**DOI:** 10.1101/2021.12.31.474593

**Authors:** Ching-Wen Chang, Krishna Mohan Parsi, Mohan Somasundaran, Emma Vanderleeden, John Cruz, Alyssa Cousineau, Ping Liu, Qi Li, Yang Wang, Rene Maehr, Jennifer P. Wang, Robert W. Finberg

**Author notes:** Those authors contributed equally to this work (Ching-Wen Chang, Krishna Mohan Parsi). Senior author (Robert W. Finberg). Corresponding author. Email addresses (C.W. Chang). Note: This paper is dedicated to the memory of Professor Robert W. Finberg. Dr. Finberg passed away unexpectedly during the preparation of the manuscript.

## Abstract

Novel pathogenic severe acute respiratory syndrome coronavirus 2 (SARS-CoV-2) continues to pose an imminent global threat since its initial outbreak in December 2019. A simple in vitro model system using cell lines highly susceptible to SARS-CoV-2 infection are critical to facilitate the study of the virus cycle and to discover effective antivirals against the virus. Human lung alveolar A549 cells are regarded as a useful and valuable model for respiratory virus infection. However, SARS-CoV-2 uses the ACE2 as receptor for viral entry and the TMPRSS2 to prime the Spike protein, both of which are negligibly expressed in A549 cells. Here, we report the generation of a robust human lung epithelial cell-based model by transducing ACE2 and TMPRSS2 into A549 cells and show that the ACE2 enriched A549^ACE2/TMPRSS2^ cells (ACE2plus) and its single-cell-derived subclone (ACE2plusC3) are highly susceptible to SARS-CoV-2 infection. These engineered ACE2plus showed higher *ACE2* and *TMPRSS2* mRNA expression levels than currently used Calu3 and commercial A549^ACE2/TMPRSS2^ cells. ACE2 and TMPRSS2 proteins were also highly and ubiquitously expressed in ACE2plusC3 cells. Additionally, antiviral drugs like Camostat mesylate, EIDD-1931, and Remdesivir strongly inhibited SARS-CoV-2 replication. Notably, multinucleated syncytia, a clinical feature commonly observed in severe COVID-19 patients was induced in ACE2plusC3 cells either by virus infection or by overexpressing the Spike proteins of different variants of SARS-CoV-2. Syncytial process was effectively blocked by the furin protease inhibitor, Decanoyl-RVKR-CMK. Taken together, we have developed a robust human A549 lung epithelial cell-based model that can be applied to probe SARS-CoV-2 replication and to facilitate the discovery of SARS-CoV-2 inhibitors.

## 1. Introduction

COVID-19 disease caused by the SARS-CoV-2 virus posed a global threat by infecting more than 250 million people and over 5 million deaths to date since its first outbreak in December 2019 (Li et al., 2021). Although SARS-CoV-2 infects various tissue types of the human body, the destruction of lung epithelial cells and the deregulated immune system led to acute respiratory distress syndrome (ARDS) and multi-organ failure in severe COVID-19 patients (Zaim et al., 2020). In addition, CoV-2 is highly contagious compared to other SARS viruses, with increased mutated variants evolving that are more transmissible and highly infectious (Zeng et al., 2022). However, the underlying molecular basis of the coronaviral disease pathogenesis remains largely unknown.

The infectivity of the SARS CoV-2 virus highly depends on the host expressing factor, angiotensin-converting enzyme 2 (ACE2). The SARS-CoV-2 spike glycoprotein protein (S) interacts with the host ACE2 receptor to initiate S1 -S2 cleavage by a cellular transmembrane serine protease, TMPRSS2. Furthermore, the endosomal proteases release the viral RNA genome into the host cell (Hoffmann et al., 2020b). Thus, the internalized viral RNA utilizes cellular protein machinery by countering the antiviral response to successfully replicating in the host cell (Hoffmann et al., 2020b). Although human stem cell-based lung epithelial CoV-2 model systems are available, need special expertise in handling cell cultures and are costly to propagate for large-scale screening applications (Huang et al., 2020). The alternate lung suitable SARS-CoV-2 permissible models, such as A549 or Calu3 cells, lack a homogenous infectious culture system with an efficient cytopathic syncytia formation, emphasizing a need for an improvised, robust platform for studying SARS viral biology and therapeutics.

In this study, we showed that A549 cells overexpressing ACE2 and TMPRSS2 stable clone(A549A549ACE2PlusC3) are highly permissible for SARS CoV-2 infection compared to Calu3 and commercial A549^ACE2+TMPRSS2^ cells. Furthermore, SARS CoV-2 infected A549ACE2PlusC cells showed cytopathic spike mediated syncytia formation, effectively blocked using Decanoyl-RVKR-CMK inhibitor. The consistent expression of SARS-CoV-2 host factors and the homogenous viral infection in A549ACE2PlusC3 cells can be an alternative improved *in vitro* disease modeling platform for drug discovery and host-viral interaction studies. Thus, A549ACE2PlusC3 cells can be a valuable resource for the current COVID-19 research community and utilized for any future challenges from SARS respiratory viruses.

## 2. Materials and methods

### 2.1. Virus, cells, plasmids, antibodies, antiviral compound

The low-passage SARS-CoV-2 (USA-WA1/2020) and icSARS-CoV-2-mNG viruses were used in this study. The USA-WA1/2020 strain (NR-52281) was provided by BEI Resources. The mNeonGreen-labeled strain was received from the World Reference Center for Emerging Viruses and Arboviruses at the University of Texas Medical Branch. Recombinant virus was prepared according to previous description (Xie et al., 2020). All work with live SARS-CoV-2 was performed in a biosafety level 3 laboratory facility (BSL3) by personnel trained to handle BSL3 agents and adhering to necessary Standard Operating Procedures.

A549 cells (CCL-185), Calu-3 (HTB-55) and Vero E6 (CRL-1586) were obtained from ATCC. Commercial cell lines, A549^ACE2^ and A549^ACE2/TMPRSS2^ were purchased from InvivoGen. 293FT was a gift from Dr. William McDougall. The cells were cultured at 37 °C in DMEM medium supplemented with 10% heat-inactivated FBS, 100 U/mL Penicillin, and 1% L-glutamine. A549 cells used for ACE2plus model establishment. Vero E6 used for virus propagation and plaque assay. 293FT was used for pseudotyped lentivirus package.

The plasmid sets for generating SARS-CoV-2 spike pseudotyped virus were provided by the BEI Resources (N-53816 and NR-53817). The generation of pseudotyped lentiviral particles was based on the protocol as previous described (Crawford et al., 2020). pcDNA3.3-WA1-SARS2, pcDNA3.3-Alpha-SARS2, pcDNA3.3-Beta-SARS2, pcDNA3.3-Delta-SARS2 were gifts from David Nemazee (Addgene plasmid #170442, #170451, #170449, 172320). pcDNA3.1-Omicron-SARS2 (# MC0101274) was purchased from GenScript. pHAGE-EF1a-ACE2PGK-puroR (internal ID 55490) and pHAGE-EF1a-TMPRSS2-PGK-puroR (internal ID 11816) plasmids were from hOrfeome 5.1 collection in the Maehr lab.

ACE2 and TMPRSS2 antibodies were purchased from R&D SYSTEMS (AF933) and Santa Cruz (sc-515727). Antiviral compounds Remdesivir and EIDD-193 were purchased from MedChemExpress (Monmouth Junction, NJ). Decanoyl-RVKR-CMK was purchased from TOCRIS (#3501). Camostat mesylate was purchased from Millipore Sigma (#SML0057). The test compounds were solubilized in DMSO to yield 10 mM stock solutions for cell culture studies.

### 2.2. Generation of A549^43.20^, A549A^CE2plus^, A549A^CE2plusC3^ cells

A549 cells were first transduced with human ACE2-expressing lentivirus (~2 × 10^5^ CFU/ml) and selected with 1ug/ml of puromycin as previous described (Koupenova et al., 2021). Next, the puromycin-resistant cells were further transduced with human TMPRSS2-expressing lentivirus (~5 × 10^5^ CFU/ml) to ensure that all cells get transduced. Over 50 single clones were selected through limiting dilution and tested for their permissiveness to SARS-CoV-2 infection. Of them, the clone 43.20 with the most efficient SARS-CoV-2 infection (~20%). To improve the infectivity, the cell populations with a higher ACE2 expression in clone 43.20 were FACS sorted using ACE2-specific antibody (AF933, R&D Systems). After puromycin selection, the infectivity significantly increased up to 40~60%. Single-cell sorting was further applied to generate ACE2plusC3 subclone that produces a homogeneous cell population. ACE2plusC3 showed stable and high levels of ACE2 and TMPRSS2 expression throughout the low- to high-passage cell culture numbers without adding puromycin in the growth medium.

### 2.3. RNA isolation and RT-qPCR

RNA was isolated from virus-containing medium or cell lysates using TRIzol LS (ThermoFisher Scientific, #10296010) according to the manufacturer’s instructions and stored at −80°C for RT-qPCR. Isolated viral RNA, its expression level was quantified using QuantiFast Pathogen RT-PCR Kit (Qiagen, #211352) and 2019-nCoV RUO Kit (IDT, #10006713). The cycling conditions were followed as the protocol recommended by the manufacturer. Isolated cellular RNA was used for cDNA synthesis using QuantiTect Reverse Transcription Kit (Qiagen, #205311). The resultant cDNAs were used to measure mRNA expression levels of ACE2 and TMPRSS2 by qPCR with gene-specific primers (human ACE2, sense 5’-GGGATCAGAGATCGGAAGAAGAAA-3’ and antisense 5’-AGGAGGTCTGAACATCATCAGTG-3’; human TMPRSS2, sense 5’-AATCGGTGTGT TCGCCTCTAC-3’ and antisense 5’ CGTAGTTCTCGTTCCAGTCGT-3’) and SYBR Green reagent. GAPDH was used as an endogenous control gene.

### 2.4. Plaque assay

Approximately 2×10^5^ Vero E6 cells were seeded to each well of 12-well plates and cultured at 37 °C, 5% CO2 for 18 h. The virus was serially diluted in MEM with 3% FBS for in vitro-generated samples, and 300 μl was transferred to the monolayers. The viruses were incubated with the cells at 37 °C with 5% CO2 for 1 h. After that, the virus-containing medium was removed and added an overlay medium to the infected cells per well. The overlay medium contained MEM with 0.42% BSA, 20mM HEPES, 0.24% NaHCO_3_ and 0.7% agarose (Oxoid, LP0028). After a 3-day incubation, plates were fixed with 4% PFA overnight and stained with crystal violet solution (Sigma-Aldrich) the next day. Plaques were counted on a lightbox.

### 2.5. Immunofluorescence staining

Cells were plated on 96-well tissue culture plates (black polystyrene microplates, Corning) and infected with the low passage of SARS-CoV2 or iscSARS-CoV2-mNG at indicated MOI and infection period. Infected cells were fixed with 4% paraformaldehyde for 30 min at room temperature, gently washed 2x in PBS, and permeabilized with 1% Triton X-100 in PBS, and blocked with 5% BSA. Fixed cells were either labeled with a human monoclonal antibody conjugated with Alexa-488 against the spike antigen (10.1371/journal.pmed.0030237) or a mouse monoclonal antibody that recognizes the NP antigen (SinoBiological, #40143-MM08) by incubation for 2 hours at 4 °C. After washing with saline-Tween 20 (0.05%), the cells were labeled with an anti-mouse goat secondary antibody conjugated with Alexa-594 by incubation for 1 h at 4 °C. To visualize the cell nuclei, the cells were counterstained with 4’,6-diamidino-2-phenylindole (DAPI) (Abcam) for 15 min at 4 °C to visualize the cell nuclei. The images were acquired with the ImageXpress Micro-XL (IXM) system by immunofluorescence with 4x or 10x objectives. The images were processed using MetaXpress Software.

For cell fusion assay, cells were plated on regular 24-well tissue culture plates for overnight to reach 90% confluence, 1ug of spike plasmid for each well was transfected into cells using transfection reagent (Mirus, TransIT-LT1) with adding DMSO or antiviral compounds for 24 hours. Transfected cells were then fixed and counterstained with DAPI. The entire plate was scanned with the Celigo Image Cytometer (Nexcelom Bioscience) and analyzed by Celigo Software.

### 2.6. Luciferase assay and Cytotoxicity

The SARS-CoV-2 spike pseudotyped virus activity was determined by bright-glo luciferase assay (Promega). The plate reader detected the luminescence two days post virus infection or without virus infection. Cell death was measured by a cytotoxicity detection kit (LDH) from Roche to assess lactate dehydrogenate activity in the culture supernatants of ACE2plus cells.

### 2.7. Statistical analyses

Data are expressed as means ± standard deviations (SD), and the significance of differences between groups was evaluated using ANOVA with Dunnett’s multiple comparisons test. All tests were performed using Prism 9 (GraphPad Software).

## 3. Results

### 3.1 Establishment of the ACE2plus cell model

To establish a robust human A549-based cell line for SARS-CoV-2 investigation, we first transduced lentiviral ACE2 and TMPRSS2 genes sequentially into A549 cells (Figure 1A).

**Fig. 1.**
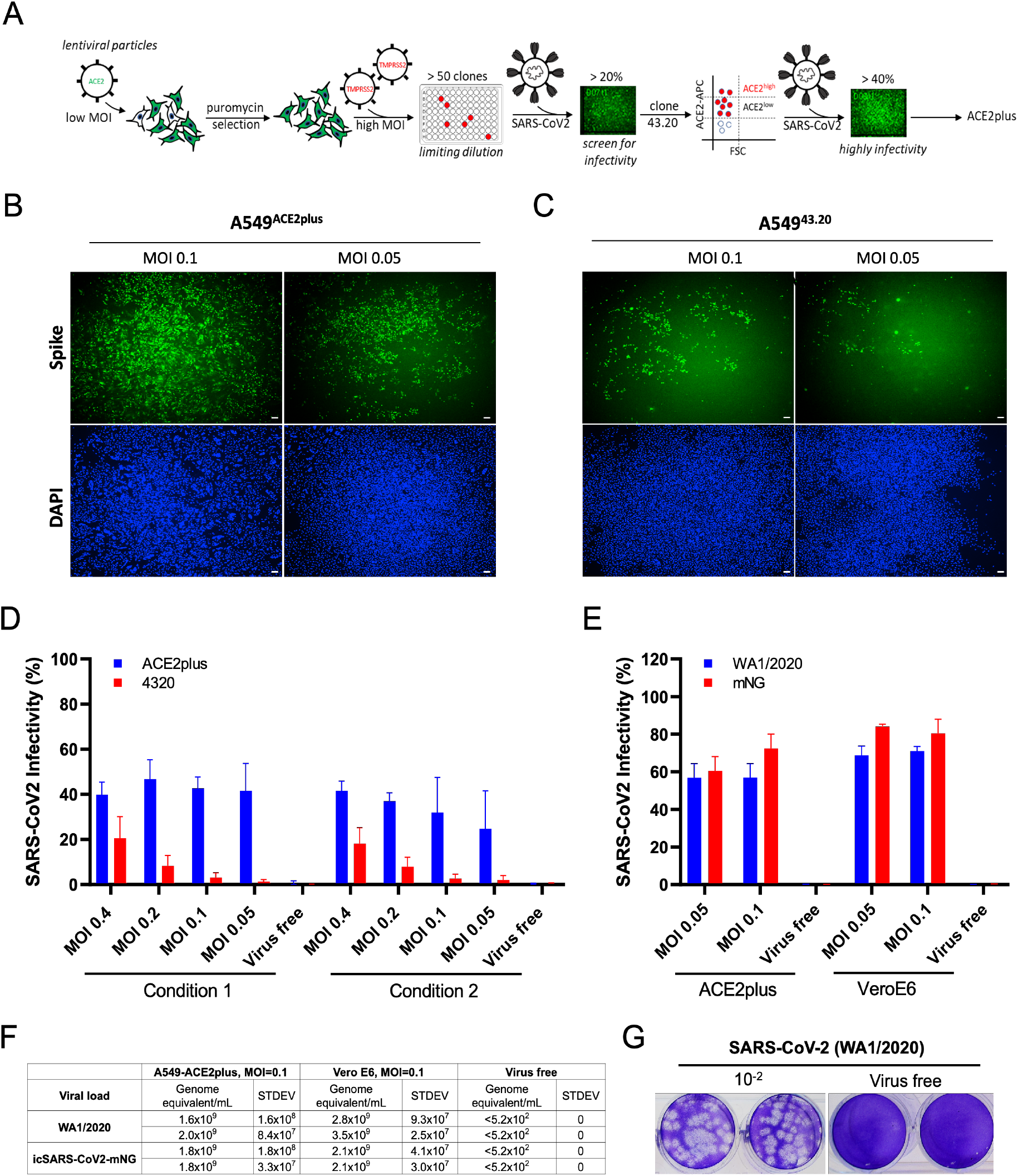
Establishment of highly permissive ACE2plus cell model for SARS-CoV-2 replication. (A) Experimental scheme to establish A549^4320^ and A549^ACE2/TMPRSS2^ (ACE2plus). SARS-CoV-2 infection observed in ACE2plus (B) and 43.20 cells (C) at 48 hours post-infection. Spike-specific antibody was used to detect infected cells; 96-well plates were imaged with the ImageXpress, 4X magnification, 100uM scale bar. (D) Scanned images were analyzed using MetaXpress software to determine the infectivity (Spike+ cells were normalized to total cell number). In condition 1, 20,000 cells per well were seeded in a 96-well plate; 15,000 cells were seeded in condition 2. (E) ACE2plus shows comparable infectivity to Vero E6 cells while challenging with SARS-CoV-2 (WA1/2020) or icSARS-CoV2-mNG (WA1/2020). (F, G) Virus-containing supernatants from infected ACE2plus cells at 48 hours post-infection were collected and analyzed by RT-qPCR and plaque assay to measure the viral load and viral particles, their ability to form plaques. The data represent the mean (±SD) from two independent experiments.

Following puromycin selection, over 50 clones were tested by SARS-CoV-2 virus infection. Clone 43.20 exhibited a higher infectivity rate of approximately 20% and was therefore selected for further ACE2 receptor enrichment by cell sorting. The sorted cell population with a high level of ACE2 expression was then referred to as ACE2plus and exhibited permissibility to SARS-CoV-2 virus replication (Figure 1B). By contrast, the parental 43.20 clone showed low levels (Figure 1C). Next, we conducted further SARS-CoV-2 infections to compare both clones with different MOIs and seeded cell numbers. Infected cells were identified by spike antibody. Images were analyzed to quantify infectivity as described in materials and methods. As Figure 1D shown, ACE2plus cells were more permissive than the parental 43.20 cells and showed higher infectivity even during low MOI virus infections, indicating that the ACE2plus model is highly susceptible to SARS-CoV-2 viral infection.

To further determine the ACE2plus model’s permissibility, we performed a comparative infection in ACE2plus and Vero E6 cells with SARS-CoV-2 virus (WA1/2020) and recombinant virus (icSARS-CoV-2mNG). Both cell types were infected with WA1/2020 (wild-type), WA1/2020 (mNG), or control growth medium (mock-infected) at low MOIs 0.05 and 0.1. Wild-type infected ACE2plus cells exhibited infection rates of approximately 60%, while Vero E6 cells showed infectivity rates around 70% (Figure 1E). ACE2plus cells exhibited a range of infectivity from approximately 60-70% across the two MOIs when infected with icSARS-CoV-2mNG, while Vero E6 cells expressed between 70-80% infectivity. To quantify viral nucleoprotein RNA levels, RT-qPCR was conducted. At 48 hours post-infection, supernatants were collected from cells treated with MOI 0.1 wild-type virus and icSARS-CoV-2-mNG, as well as mock-infected, and processed for RNA extraction. Vero E6 cells showed 1.75 fold higher infectivity than ACE2plus when treated with wild-type virus and approximately 1.2 fold higher infectivity than ACE2plus when challenged with icSARS-CoV-2-mNG (Figure 1F). Released virus particles from infected ACE2plus cells were also tested using plaque assay to determinate the virus activity (Figure 1G). Altogether, those results demonstrate the comparable efficacy of ACE2plus and Vero E6 cells as models for SARS-CoV-2 infections and shows that ACE2plus is a viable human cell model.

### 3.2 Characterization of the ACE2plus cell model in comparison to commercial A549^ACE2/TMPRSS2^ cells

ACE2 and TMPRSS2 are key receptors for SARS-CoV-2 entry. To measure levels of *ACE2* and *TMPRSS2* mRNA expression in the ACE2plus model, RT-qPCR was performed on supernatants collected from ACE2plus, parental A549, Calu3 and two commercial A549^ACE2^ (IVG-A), A549^ACE2/TMPRSS2^ cell lines (IVG-AT). Relative mRNA expression was normalized to housekeeping gene GAPDH. *ACE2* mRNA expression levels in ACE2plus and IVG-AT cells were similar (Figure 2A). However, ACE2plus cells expressed a higher level of *TMPRSS2* mRNA than IVG-AT and Calu3. Both gene expressions in parental A549 are extremely low. Next, we used flow cytometry on ACE2plus and IVG-AT cells to determine the cell-surface ACE2 protein expression. As expected, the ACE2plus cell population showed > 95% expression of ACE2 while the IVG-AT cells only exhibited 33% expression (Figure 2B). Although the ACE2 positive population can be increased after drugs selection, the IVG-AT cells grow very slowly. Thus, cell growth rates were determined by staining ACE2plus and IVG-AT cultures with DAPI using Celigo imaging and software (Figure 2C) to determine cell count in 24-hour time increments over a total period of 96 hours. Both cell lines were seeded in a 96-well plate at 1×10^4^ cells per well. After 24-hour, the ACE2plus cell grows faster and more consistent than the commercial IVG-AT cell line.

**Fig. 2.**
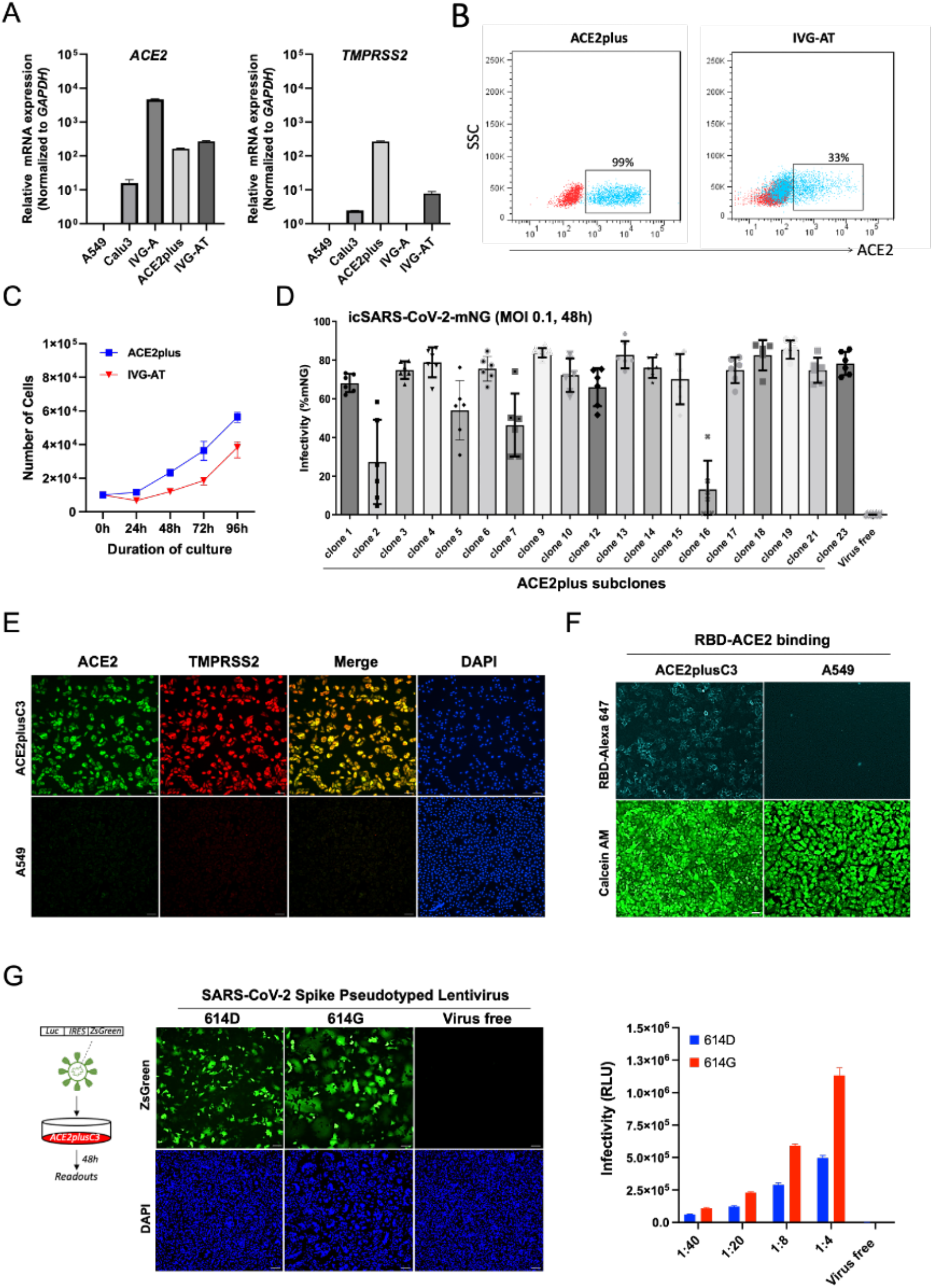
Characterization of ACE2plus cell model and its subclone ACE2plusC3. (A) The mRNA expression levels of ACE2 and TMPRSS2 in indicated cell lines were measured by RT-qPCR. A549^ACE2^ (IVG-A) and A549^ACE2/TMPRSS2^ (IVG-A/T) are commercial cell lines used as a control (B). The cell surface ACE2 expression level was measured by flow cytometry using live cells. (C) Cell number at different time points was evaluated by DAPI counting using Celigo Image Cytometer. (D) Single-cell derived clones were generated through FACS sorting using ACE2-specific antibody. Following expansion, clones derived from ACE2plus cells were infected with icSARS-CoV2-mNG virus to determine their infectivity. (E) The expression of ACE2 and TMPRSS2 proteins in clone 3 (ACE2plusC3) was examined by immunofluorescence microscopy (10x magnification). (F) RBD/ACE2 binding in living cells. Recombinant SARS-CoV-2 Spike RBD proteins were labeled using Alexa Fluo 647 Labeling Kit (ThermoFisher Scientific) and incubated with cells on ice for 30 minutes. RBD-Alexa 647 signal was detected by Celigo Image Cytometer. Calcein AM was used to detect living cells. (G) ACE2plusC3 cells are infectable with SARS-CoV-2 Spike-pseudotyped lentivirus (PV). Microscope images of ZsGreen expression in ACE2plusC3 at 48 hours post-incubation with 614D or 614G PV. A series of diluted virus was applied, and infectivity was measured via relative luciferase units (RLU). Each data represents the mean and standard deviation. 100uM scale bar.

Considering cell heterogeneity, we further optimized the ACE2plus model by single-cell sorting. To do this, the ACE2plus cell was incubated with an ACE2-specific antibody for Fluorescence Activated Cell Sorting (FACS). We then successfully expanded 23 clones and challenged them with icSARS-CoV-2-mNG virus to compare the infectivity of each clone (Figure 2D). Of them, clone 3 (ACE2plusC3) was then selected due to its high expression of ACE2 and less variation among infected cells (data not shown). We then used Immunofluorescence staining for ACE2 and TMPRSS2 in ACE2plusC3 cells. A549 was used as a staining control. A549 cells exhibited negligible ACE2 and TMPRSS2, while ACE2plusC3 showed strong and ubiquitous expression for both (Figure 2E). Next, we examined if the ACE2 receptor on the ACE2plusC3 cell surface can be recognized by SARS-CoV-2 Spike-RBD protein. As Figure 2 F shows, recombinant RBD proteins can specifically bind to the ACE2 receptor and be internalized rapidly within 45 minutes (data not shown). According to recent reports, spike D614G mutation is associated with ACE2 receptor binding and results in an increase in infectivity of the SARS-CoV-2 (Cheng et al.). To test this, the 614D and 614G of Spike-pseudotyped lentiviral particles were prepared and used for infections. As *luciferase* and *ZsGreen* genes were designed as reporters in the system, the infectivity can be easily measured by ZsGreen protein expression or luciferase assay (Crawford et al., 2020). As a result, 614G showed stronger infectivity than 614D in a dose-dependent manner (Figure 2G). This data supports the use of ACE2plusC3 for SARS-CoV-2 lentiviral infection assays. Taken together, those results provide evidence that the ACE2plus cell line is an ideal model for the SARS-CoV-2 study.

### 3.3 SARS-CoV-2-Spike-mediated cell-cell fusion

Cell-cell fusion allows viruses to infect neighboring cells, and it was recently discovered that the polybasic S1/S2 site of SARS-CoV-2 Spike is required for efficient infection of human lung-derived cells and promotes syncytium formation (Cheng et al., 2020; Hoffmann et al., 2020a). Thus, it might be essential to understand the ability of syncytium formation between SARS-CoV-2 spike variants, as the large size of syncytia is reported to constitute a hallmark of COVID-19-associated pathology (Bussani et al., 2020). To address this, we first established a mCherry stable cell line using the ACE2plusC3 model. After sorting, over 98% of cells express mCherry protein, and this cell model can be applied to real-time observe the kinetic of spike-mediated cell-cell fusion without no need to coculture with other cells. Next, we transfected an equal amount of indicated spike plasmids into ACE2plusC3-mCherry cells for 24 hours. WA1/2020 spike used as a reference, and pCDNA empty vector acts as a negative control. To visualize and capture whole well images, an entire 24-well plate was scanned using Celigo Image Cytometer (Figure 3A). As a result, delta and beta spikes clearly induced giant syncytia formation (> 300uM) with multinucleated enlarged cells compared with WA1/2020 or alpha spike (Figure 3B). The size and number of syncytia was dramatically reduced while adding CMK or Camostat inhibitors to transfected cells. Notably, omicron spike seems to poorly induce cell fusion in ACE2plus-mCherry cells. Those results indicate that the ACE2plusC2 model can be used to study spike-mediated cell fusion.

**Fig. 3.**
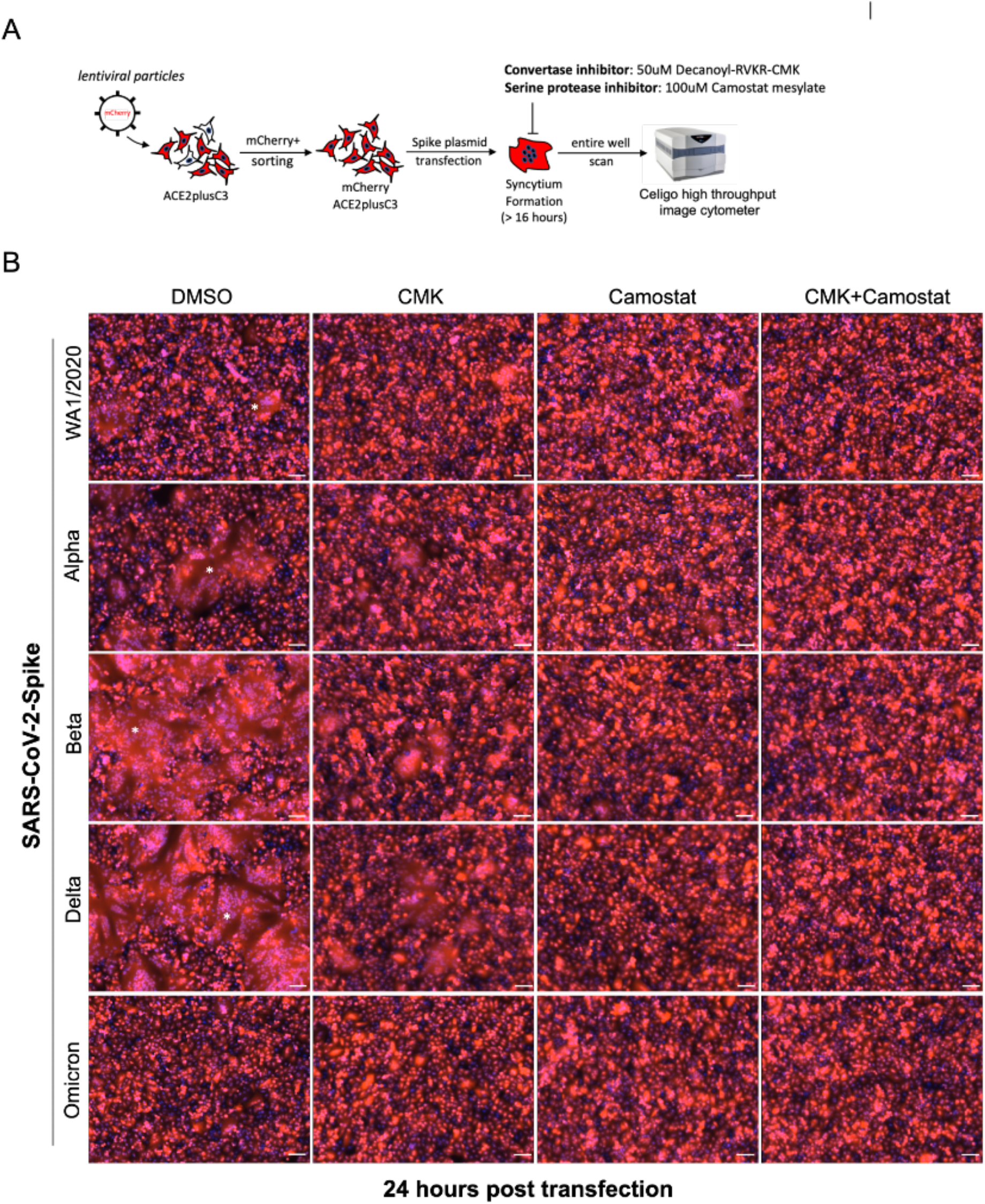
SARS-CoV-2-Spike-mediated syncytium formation in ACE2plusC3 cells. (A) Experimental scheme to observe syncytium formation. To visualize and capture whole well images, an entire 24-well plate was scanned (B) Representative images from two independent experiments. SARS-CoV-2 Spike plasmids were transfected into cells with or without protease inhibitor treatment. After 24 hours, cells were fixed and scanned using Celigo Image Cytometer. mCherry and DAPI images were merged by Celigo Software. 100uM scale bar. Star shows the area of the syncytium.

### 3.4 Evaluation of the antivirals against SARS-CoV-2 using ACE2plusC3 cell culture model

To test the utility of the ACE2plus model in anti-viral drug screening, we evaluated the efficacy of Camostat mesylate, EIDD-1931, and Remdesivir in inhibiting SARS-CoV-2 infection. ACE2plusC3 cells were pretreated with drugs for one hour prior to infection. Then cells were infected with icSARS-CoV-2-mNG at MOI 0.1. After 48 hours, supernatants were collected for cytotoxicity assay, infected cells were fixed and quantified by IXM image system. Clearly, even at low concentration of EIDD-1931 (1uM) and Remdesivir (0.1uM) we observed approximately 50% reduction of NP fluorescence (Figure 4). EIDD-1931 and Remdesivir, which target virus RNA-dependent RNA polymerase, have been reported to be potent antivirals against SARS-CoV-2 (Miller et al., 2021). Camostat mesylate is regarded as an antiviral agent, as it inhibits many of the serine proteases that SARS-CoV and SARS-CoV-2 use for virus-to-host cell membrane fusion, like TMPRSS2, and TMPRSS11 (Breining et al., 2021). As a result, treating cells with 100uM of Camostat mesylate resulted in significant inhibition (Figure 4). This is consistent with recent reports that Camostat mesylate significantly reduced SARS-CoV-2-driven entry and infection in lung cell line Calu-3 and primary human lung cell (Hoffmann et al., 2020b).

**Fig. 4.**
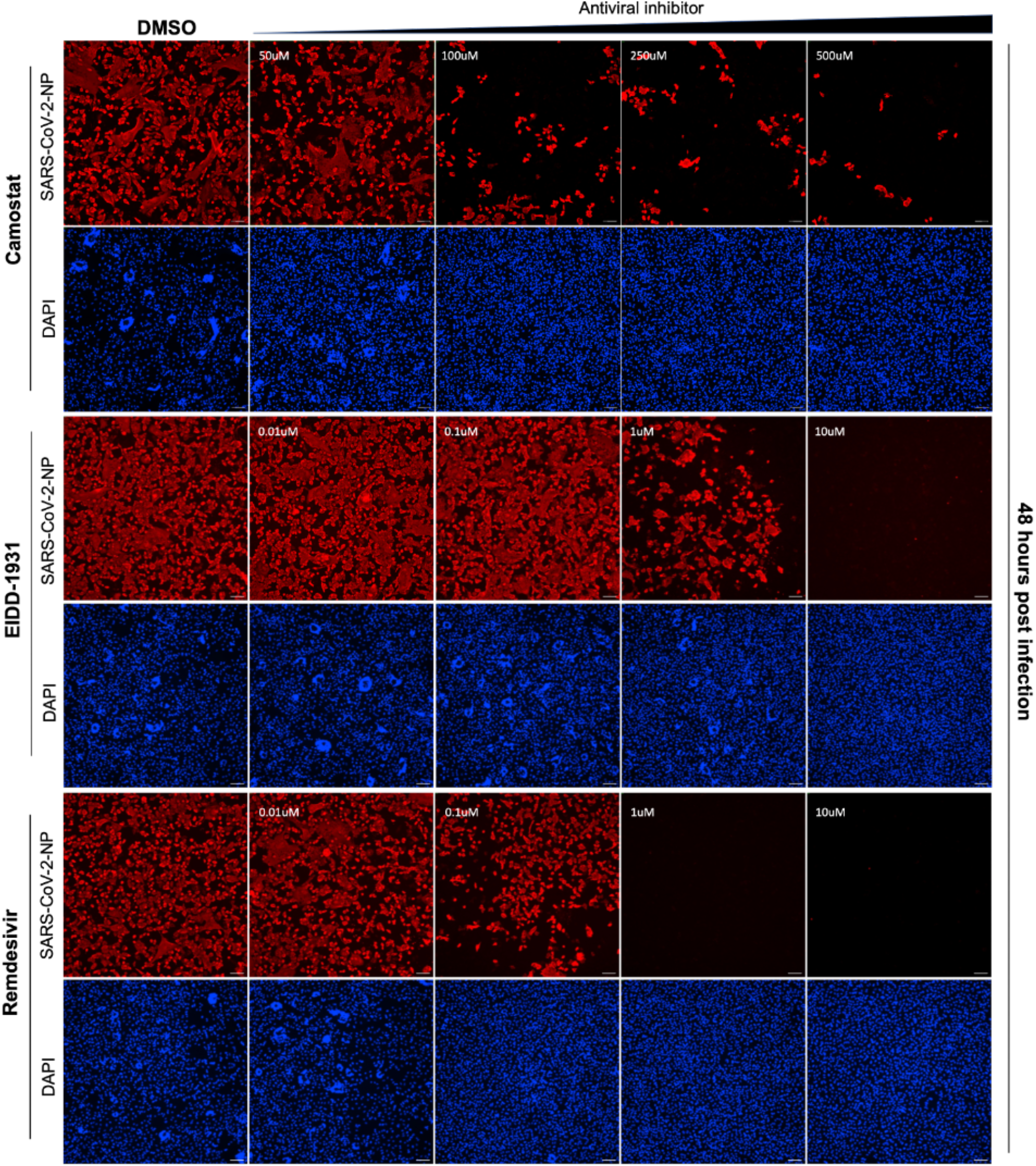
Dose-dependent inhibition in SARS-CoV-2 infection. Representative images from two independent experiments. Cells were plated in a 96-well plate overnight as described in materials and methods. The next day, cells were infected with icSARS-CoV-2-mNG at MOI 0.1 and treated with a range of concentrations of antiviral drugs for 48 hours. After that, cells were fixed and stained with anti-NP antibody, followed by secondary antibody incubation (conjugated with Alexa-594). DAPI was used for nuclear counterstain. Images were scanned by ImageXpress using 10x magnification, 100uM scale bar.

To test potential dose-dependent antiviral activity of those drugs in our cell model, we treated ACE2plusC3 with different concentrations of those drugs and infected the cells with icSARS-CoV-2-mNG at MOI 0.1. Cells were fixed at 48 hours post-infection and determined the infectivity as abovementioned. To quantify the inhibition of those drugs, the 48 hour time point values from two independent experiments (n=6) were plotted on a semi-logarithmic graph to establish the half-maximal inhibitory concentration value. Camostat mesylate, EIDD-1931, and Remdesivir exhibited potent antiviral effect with IC_50_=59.98 uM, 0.84uM, and 0.14uM, respectively (Figure 5). No apparent cytotoxic effect was observed in cells. These results demonstrated that our ACE2plucC3 cell model can be applied for evaluation of antiviral drugs and might be potentially developed for high-throughput screening.

**Fig. 5.**
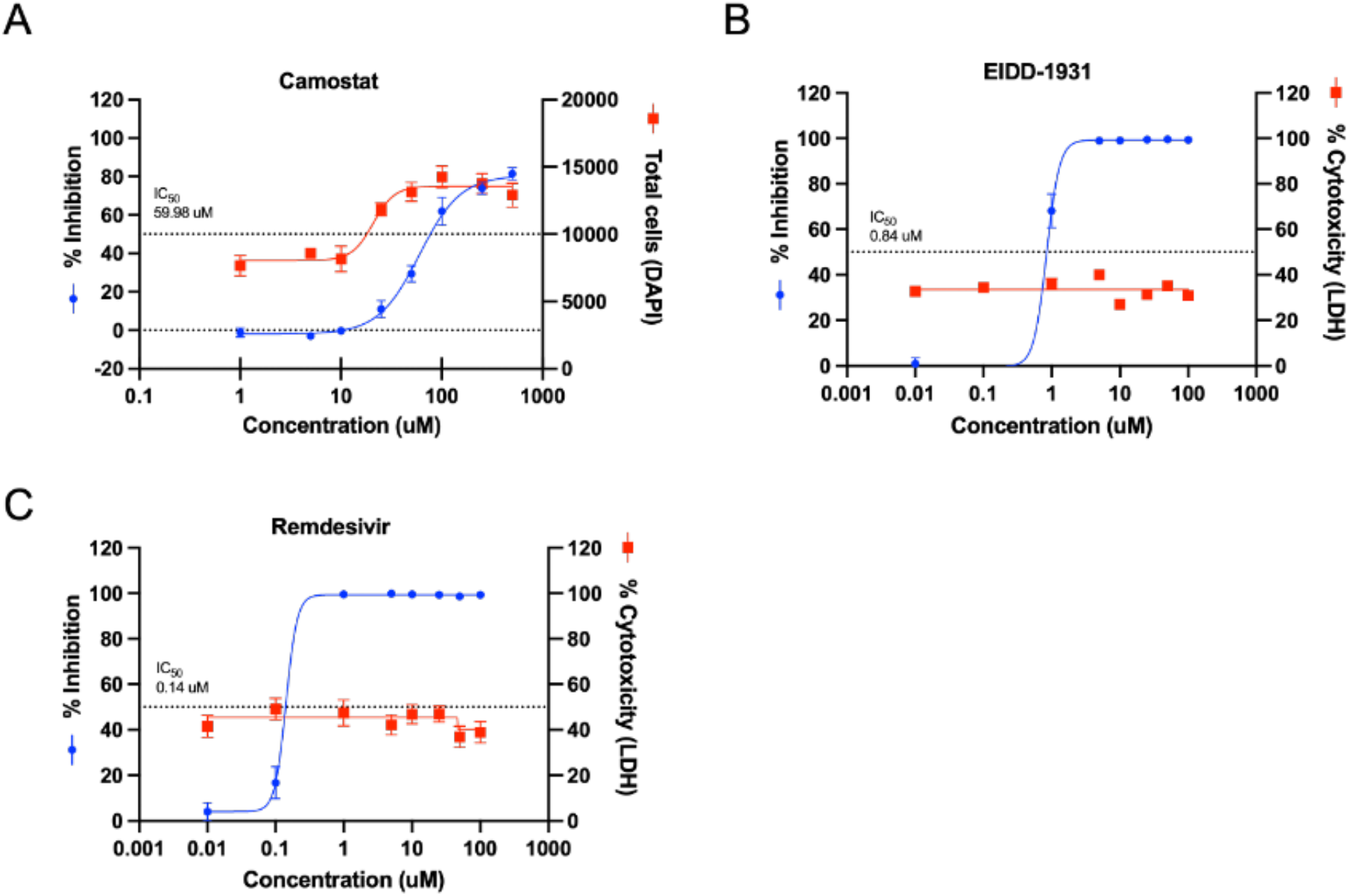
Antiviral efficacy of Camostat, Remdesivir and EIDD1931 in ACE2plusC3. Cells were infected with icSARS-CoV-2-mNG at MOI 0.1 and treated with a range of concentrations of antiviral drugs for 48 hours. Supernatants were collected for LDH cytotoxicity assay. Fixed cells were stained with anti-NP antibody, followed by secondary antibody incubation (conjugated with Alexa-594). DAPI was used for counterstain. The images were acquired with the ImageXpress system by immunofluorescence with 4x and processed by MetaXpress Software to calculate the infectivity. GraphPad was used to draw the dose-response curve and determine the IC_50_. The data represent the mean (±SD) from two independent experiments.

### 3.5 Decanoyl-RVKR-CMK inhibits SARS-CoV-2 infection in ACE2plusC3 cells through suppressing spike-mediated cell fusion

In the transfection experiments, we found that the spike-mediated cell fusion was reduced by treatment with furin inhibitor Decanoyl-RVKR-CMK (Figure 3B). Because several reports have demonstrated that the cleavage of the SARS-CoV-2 spike protein at a putative furin cleavage site (RRARS) at R685/S686 is critical for spike-mediated cell-cell fusion (Cheng et al.; Cheng et al., 2020; Hoffmann et al., 2020a). We also observed extensive syncytial phenotype in SARS-CoV-2-infected ACE2plus cells. We next investigated the efficacy of the furin inhibitor Decanoyl-RVKR-CMK to determine whether the furin protease is required for syncytium formation in ACE2plucC3 cells. To do this, cells were infected with SARS-CoV-2 (WA1/2020) at MOI 0.1 and inoculated with a range of concentrations of Decanoyl-RVKR-CMK for 36 hours before staining for DAPI and nucleocapsid protein expression. Mock-treated cells were infected with the virus and received no drug inoculation. In a concentration-dependent manner, cytotoxicity effects were not observed. (Figure 6D). As a result, syncytia formation were markedly observed in virus-infected cells, and the syncytial phenotype and infected cells were less clearly prominent in the presence of the CMK furin inhibitor (Figure 6A, 6B). To quantify the syncytia number, each image captured from 4x object was analyzed based on the size (> 100uM) and the number of nuclei (> 5). As Fig 6C shown, the syncytia number is significantly reduced by CMK inhibitor. To further test inhibition of viral infection in these samples, plaque assays were performed on Vero E6 cells with supernatants collected at 36 hours post-infection. The virus activity was inhibited in CMK-treated samples (Figure 6E). Besides, mock-treated samples had a virus titer of approximately 1.75×10^5^ plaque-forming units per mL (PFU/mL), and decreased to 0.8×10^5^ PFU/mL in the 100uM CMK-treated samples (Figure 6F). Thus, the furin-dependent process possibly contributes to syncytium formation in ACE2plusC3 cells.

**Fig. 6.**
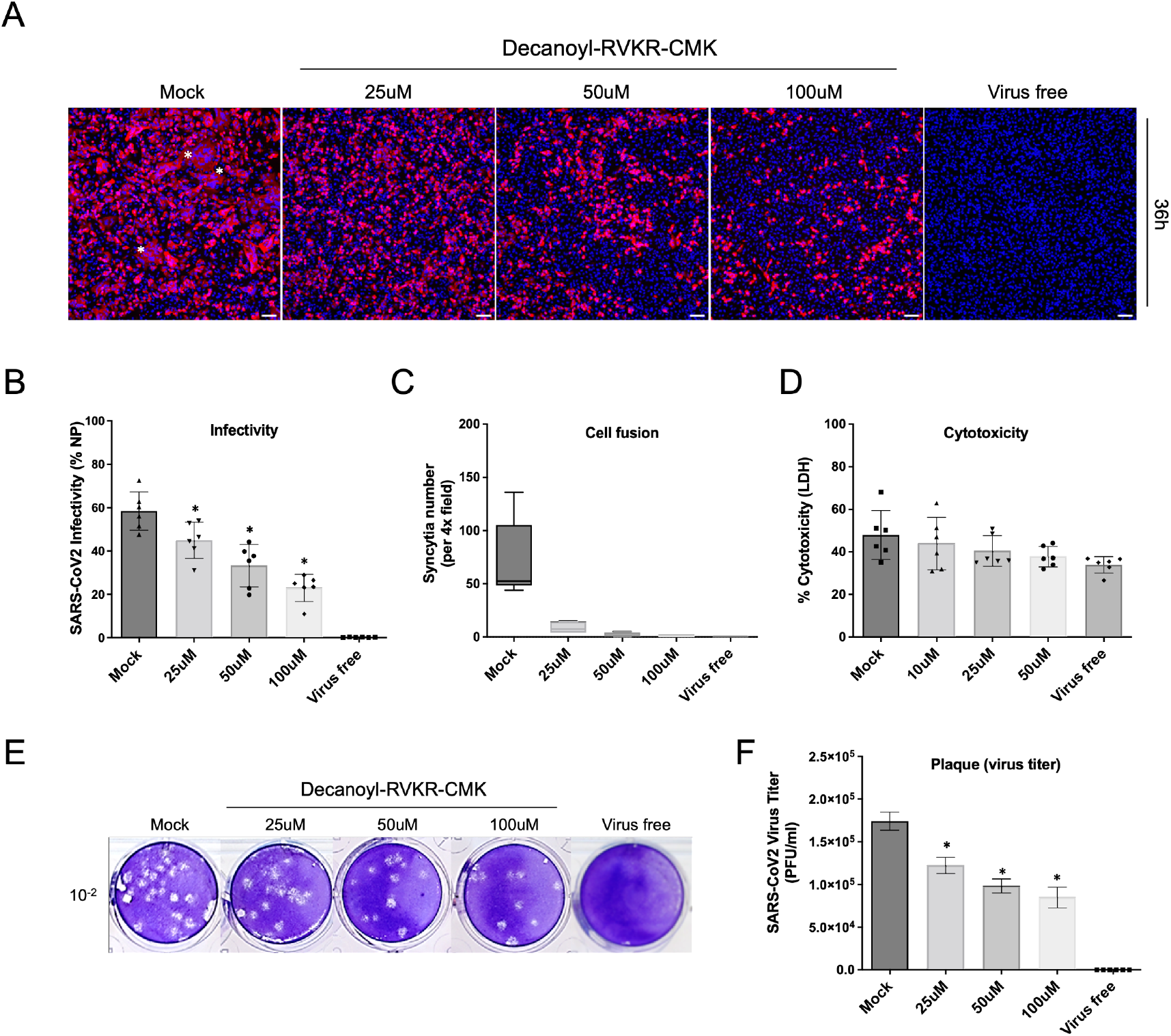
Decanoyl-RVKR-CMK inhibits SARS-CoV2 infectivity by suppressing Spike-mediated cell fusion in ACE2plusC3. (A) Microscope images showing viral nucleocapsid protein expression (red) in infected cells, DAPI (blue) was used for nuclear counterstain. Images were scanned by ImageXpress using 10x magnification. Cells were infected with WA1/2020 strain at an MOI of 0.1 in the presence of furin inhibitor drug dilutions for 36 hours. (B, C) Quantifying the virus infectivity and syncytia number via ImageXpress, Syncytia were determined based on the size (> 100uM) and the number of nuclei (> 5). (D) Cytotoxicity was measured by LDH assay. (E) Plaque assay. Supernatants were collected at the 36-hour timepoint post-infection. (F) Virus titer was quantified from plaque numbers. Data are representative of the mean and SEM of two independent experiments (P* < 0.05). Star shows the area of the syncytium.

## 4. Discussion

The current pandemic caused by SARS-CoV2 is completing its second year of global devastation of human lives. While novel vaccine strategies have provided imminent protection and slowed the pace of spreading infection, this approach has been seriously challenged by newly emerging variants. There is critical need for effective antivirals along with the current vaccine strategy to successfully combat this as well as newly emerging pandemics caused by respiratory viruses. A reliable cell model that reproduces the SARS-CoV-2 life cycle is therefore required to help us better understand the virus-host interactions and to discover novel antiviral drugs. Studies have shown that ACE2 receptor is considered essential for SARS-CoV-2 entry, and the serine protease TMPRSS2 for spike protein priming and spike-mediated cell fusion (Hoffmann et al., 2020b). Since the outbreak of the COVID-19 pandemic, many efforts have been focused on to establish various cell models to perform relevant in vitro studies. Several commonly used cell lines such as 293^ACE2^ and Vero E6 have been widely utilized for studying SARS-CoV-2 virus entry, replication, and antivirals (Cheng et al., 2020; Hoffmann et al., 2020b; Zeng et al., 2022). However, they may not be suitable cell models to investigate the pathological mechanism of the host cell in response to the virus infection, as they were not derived from human lung tissue and lack cytopathic effect (CPE) as well as type I interferon genes expression (Osada et al., 2014). To address this, we systematically developed a highly permissive human lung-based ACE2plus cell model which originated from human lung epithelial cell line A549. By comparing to other cell lines used for susceptibility to SARS-CoV2 infection and replication, ACE2plus expressed a higher level of *ACE2* and *TMPRSS2* than Calu3 and IVG-AT (A549^ACE2/TMPRSS2^, InvivoGen) and showed a faster and more consistent growth rate than the commercial IVG-AT cell line, indicating that it is an easily manipulated cell model for large-scale screening applications.

As ACE2plusC3 was derived from a single cell colony, with homogenous population, it is ubiquitously expressed ACE2 and TMPRSS2 proteins. In this study, we evaluated the longevity of the ACE2plusC3 model’s susceptibility to SARS-CoV-2 infection using passage 16~18 of cells, they exhibited similar infectivity levels of 70~80% as early passage of ACE2plusC3 cells. We also evaluated the sensitivity of ACE2plusC3 model for SARS-CoV-2 antivirals using Camostat mesylate and two FDA-approved drugs, Remdesivir and EIDD-1931(molnupiravir’s active metabolite), our results showed potent antiviral effect with IC50=59.98 uM, IC_50_=0.14uM, and IC_50_=0.84uM, respectively. We noticed that there is a comparative data of the Remdesivir IC_50_ in different cell lines. For example, the reference drug Remdesivir has been showed differences in IC_50_ in between Vero (IC_50_=10uM) and Calu-3 (IC_50_=1.3 uM) cells (Jang et al., 2021; Ko et al., 2021). This indicates that our ACE2plusC3 is a more sensitive cell model and better than Vero cells for developing an antiviral screening assay.

Furthermore, we clearly observed the syncytium formation in cells infected with wild type WA01/2020 strain. Syncytia were also evident in ACE2plusC3 cells transduced with different variants of Spike proteins. We found that the beta- and delta-Spike both had stronger fusogenic activity than WA1/2020 and others. Notably, extensive cell fusion caused by the delta Spike led to giant syncytium formation and induce cell death. By contrast, cell-cell fusion was barely observed with overexpressed omicron Spike at the same condition. Our results are in line with recent reports that show delta-Spike possess higher fusion activity and syncytium formation ability than the parent WA1/2020, and thus likely induced cell-cell fusion in the respiratory tract to cause severe pathogenicity reported in infected individuals (Mlcochova et al., 2021). Although very little is understood about the omicron variant, the severity of disease is reportedly much less than the delta variant. Our results strongly suggested that the Omicron spike protein induced relatively poor cell fusion similar to recent reports in 293^ACE2^ and VeroE6^TMPRSS2^ cells (Meng et al., 2021; Zhao et al., 2021).

Coexpression of ACE2 and TMPRSS2 strongly correlates with the virus susceptibility and cell-to-cell infection. In addition to the Spike protein inducing cell-cell fusion, furin and TMPRSS2 play important roles in this process of spike-mediated cell fusion (Hoffmann et al., 2020b). In this report, using furin convertase inhibitor, we have provided evidence that the relatively robust cell infection efficiency of SARS-CoV-2 in ACE2plus is most likely dependent on its higher cell fusion capability (compared to the DMSO treatment). However, we cannot exclude the possibility that other molecules are involved in viral recognition and entry. Further investigation is warranted to identify and tease out the exact roles played by these factors. Overall, we have established a robust human lung-based cell model for SARS-CoV-2 infection. Our data on SARS-CoV-2 virus production, pseudotyped virus infection, Spike-mediated cell fusion, and antiviral test highlight the importance of our cell model, which might provide as a powerful tool to facilitate the study of the emerging SARS-CoV-2 variants.

## Acknowledgements

This work was supported by the Department of Defense (DoD) COVID-19 Expansion Award W81XWH2110029 to Dr. Robert W. Finberg. The following reagent was obtained through BEI Resources, NIAID, NIH: SARS-Related Coronavirus 2, Wuhan-Hu-1 Spike-Pseudotyped Lentiviral Kit, NR-53816 and NR-53817

## Contributions

C.C. designed research; C.C., K.M.P., E.V., M.S., P.L., and J.C. performed the experiments; Q. L. and Y. W. prepared anti-spike antibody and tested its specificity. C.C., R.W.F, J.W., R.M, M. S. analyzed the data; C.C., K.M.P. and E.V. wrote the manuscript.

